# SAPdb: A database of nanostructures formed by self-assembly of short peptides

**DOI:** 10.1101/685149

**Authors:** Deepika Mathur, Harpreet Kaur, Anjali Dhall, Neelam Sharma, Gajendra P. S. Raghava

## Abstract

**Background:** Nanostructures generated by self-assembly of peptides yield nanomaterial that has many therapeutic applications, including drug delivery and biomedical engineering, due to their low cytotoxicity and higher uptake by targeted cells owing to their high affinity and specificity towards cell surface receptors. Despite the promising implications of this rapidly expanding field, there is no dedicated resource to study peptide nanostructures.

**Result:** This study endeavours to create a dedicated repository of short peptides, which may prove to be the best models to study ordered nanostructures formed by peptide self-assembly. SAPdb has a repertoire of 1,049 entries of experimentally validated nanostructures formed by the self-assembling of small peptides. It includes 701 entries are of dipeptides, 328 entries belong to tripeptides, and 20 entries of single amino acid with some conjugated partners. Each entry encompasses comprehensive information about the peptide such as chemical modifications in the peptide sequences, the type of nanostructure formed, and experimental conditions like pH, temperature, and solvent required for the self-assembly of the peptide, etc. Further, our analysis has shown that the occurrence of aromatic amino acids favours the formation of self-assembling nanostructures, as indicated by a large number of entries in SAPdb contain aromatics amino acids. Besides, we have observed that these peptides form different nanostructures under different experimental conditions. SAPdb provides this comprehensive information in a hassle-free tabulated manner at a glance. User-friendly browsing, searching, and analysis modules are integrated for easy retrieval and comparison of data and examination of properties. We anticipate SAPdb to be a valuable repository for researchers engaged in the burgeoning arena of nanobiotechnology.

**Availability:** The database can be accessed on the web at https://webs.iiitd.edu.in/raghava/sapdb.

## Introduction

Peptides have been reported as key players in diverse fields like immunotherapeutic [1–7], disease biomarkers [8–10], antibacterial [11–15], antiviral [16–20], anticancer [21–25], antiparasitic [26– 30], antihypertensive [31–33] drugs owing to their properties such as cell-penetration [34], stability [35–37] and low toxicity [38, 39]. Besides these areas, peptides are rapidly gaining the attention of researchers in the field of nanobiotechnology [40–43] by virtue of their property to self-assembly into well-defined nanostructures. Advantages of self-assembled short peptides for use as nanomaterial relative to conventional materials include simple structure, fast and low-cost synthesis, better chemical and physical stability, diversity in morphology, ease of synthesis in large quantities. In addition, biocompatibility, biodegradability, low cytotoxicity, and higher uptake by targeted cells of SAPs (self-assembling peptides) play a significant role in their therapeutic applications [44, 45]. Owing to these attractive properties, bioactive peptides with the ability to undergo self-assembly are being explored to serve as building blocks of hydrogels and scaffolds in cell culture [46] and tissue engineering [47, 48], controlled drug delivery in response to changes in pH [49, 50] for diagnostics and biosensors [51], as well as in the field of bioelectronics [52] and material sciences [52].

Self-assembly of peptides can lead to the formation of well-defined nanostructures like nanofibers, nanorods, nanoparticles, hydrogels, nanotubes, etc. [53–61]. The assembly of these nanostructures depends on weak non-covalent interactions like Van der Waals force, hydrophobic interactions, hydrogen bonds, and π-π stacking [62–65]. The lack of a comprehensive understanding of the properties and mechanisms of the self-assembly of peptide nanostructure formation affects their configuration and utility. Thus, it is a challenge to control the shape, size, and stability of these nanostructures during synthesis for successful integration and translation into biosensors and controlled release devices for drug delivery.

Though various studies have been carried out to design peptide-based nanoparticles, to the best of the authors’ knowledge, there is no dedicated platform that maintains comprehensive information about nanoparticles formed by the self-assembly of peptides. A systematic collection and compilation of experimental data is needed to examine the mechanisms and interactions governing the self-assembly of peptides into nanostructures for facilitating the rational designing of morphologies and the size of peptide assemblies. The present report is the first attempt to develop a repository of short peptides that undergo self-assembly to form nanostructures. Dipeptides and tripeptides are the smallest of known peptide self-assemblers, which show fascinating morphologies and functionalities besides being cost-efficient and fast to synthesize. “SAPdb” database of such short peptides will be very beneficial to study how experimental conditions and chemical modifications in the amino acid sequence of peptides affect the bottom-up process of self-assembly to form well-defined ordered nanostructures.

## Methods

### Data Collection

To collate the relevant information for di- and tri-peptides that self-assemble to form discrete and ordered nanostructure, PubMed was queried to obtain the research articles. Keywords like “(tripeptide AND self-assembly)”, “(tripeptide AND nanostructure)”, “(tripeptide AND nanotube)”, “(tripeptide AND nanofiber)”, “(tripeptide AND nanorod)”, “(tripeptide AND hydrogel)”, “(tripeptide AND nanosphere)” and “(tripeptide AND nanoparticle)” for self-assembling tripeptides, while to collect research articles relevant to self-assembling dipeptides keywords “(dipeptide AND self-assembly)”, “(dipeptide AND nanotube)”, “(dipeptide AND nanostructure)”, “(dipeptide AND nanorod)”, “(dipeptide AND nanosphere)”, “(dipeptide AND nanofiber)”, “(dipeptide AND hydrogel)”, “(dipeptide AND nanoparticle)” were used collected around ∼1500 publications till July 2019. We screened the articles obtained and included only those for further data curation, where information about peptides forming self-assembled nanostructures was available. These self-assembling peptides form various nanostructures, i.e., nanosphere, nanotube, hydrogel, nanovesicle, nanofibers, etc. as represented in Figure 1.

**Figure 1:**
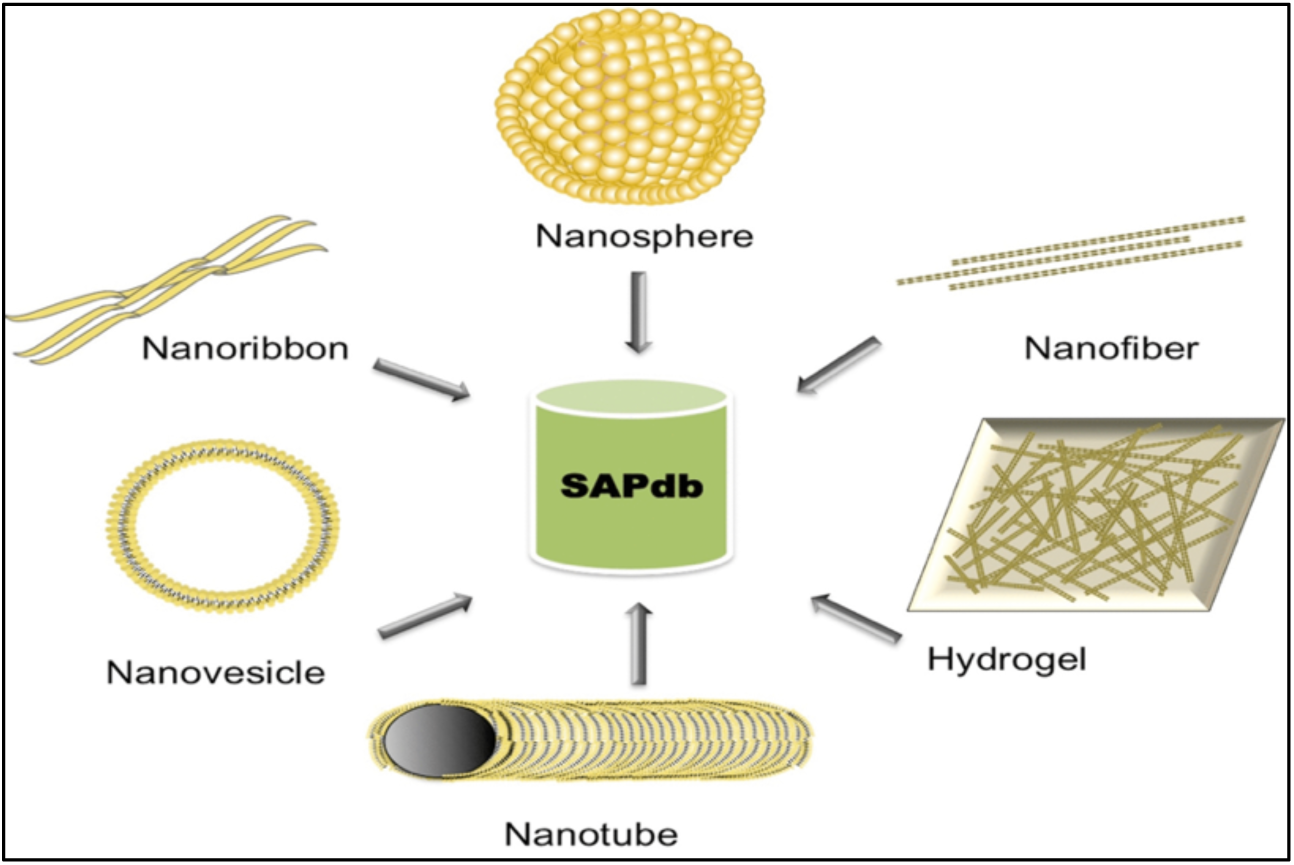
Description of the formation of different nanostructures from the self-assembly of peptides. After the thorough screening, relevant information extracted from selected articles concerning the sequence of peptides, N-terminal modifications, C-terminal modification, a technique to analyze self-assembled structure, method, type of self-assembly, size of the self-assembled structure, and the conditions under which peptide formed the self-assembled structure such as solvent, concentration, temperature, pH and incubation time, etc. To provide information regarding the effect of experimental conditions on the self-assembly of peptides like concentration, solvent, pH, incubation time, etc. on the self-assembly of peptides, we made multiple entries of the same peptide if it was reported under different experimental conditions. This extensive information is systematically cataloged in a tabulated manner. Consequently, 1,049 entries were collated in the SAPdb database from 301 research articles.

### Architecture and web - interface of the Database

SAPdb was designed using Apache HTTP Server (version 2.2.17) on the Linux system following the collection and compilation of significant information. SAPdb is based on MySQL at the back end that was implemented to maintain the information and HTML5, PHP, and JavaScript. They were deployed to execute the front end in order to build a mobile, tablet, and desktop compatible web-resource. Different modules were integrated into SAPdb for data compilation, retrieval, and exploration. The complete architecture depicting information and tools embedded into SAPdb represented in Figure 2.

**Figure 2:**
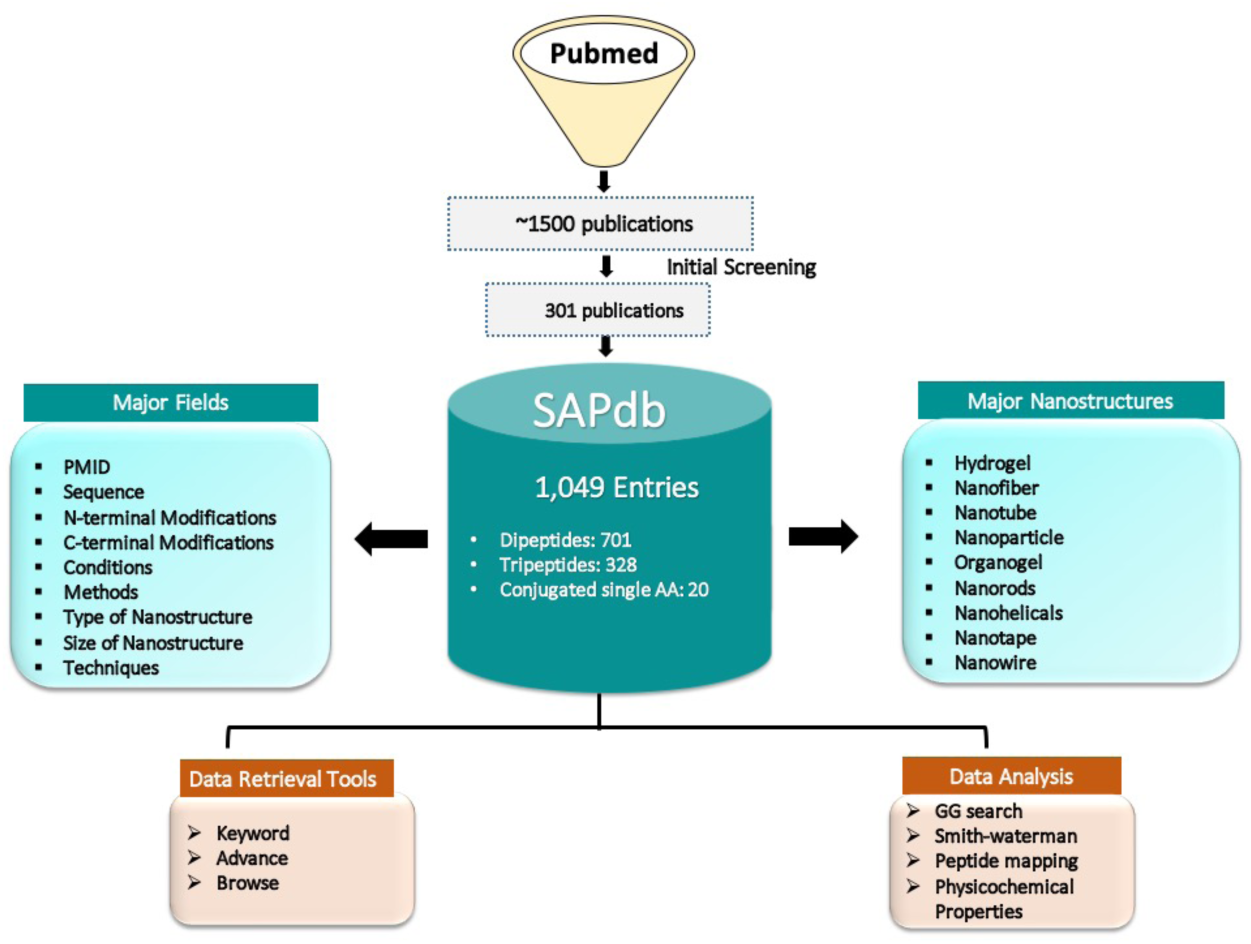
Architecture depicting the information and tools integrated in SAPdb.

### Organization of Database

In SAPdb, collated information was classified into primary and secondary information. The primary information includes fields like (i) PubMed ID, (ii) peptide sequence, (iii) N-terminal modification, (iv) C-terminal modification, (v) non-terminal modification, (vi) technique, (vii) method, (viii) solvent, (ix) concentration of peptide, (x) pH, (xi) temperature, (xii) incubation time, (xiii) type of self-assembly, (xiv) size of self-assembled structure and (xv) stability of self-assembled structure. While in secondary information (xvi) SMILES and (xvii) tertiary structure of peptides were included.

## Results

### Statistical analysis of data of SAPdb

SAPdb is a collection of 1,049 entries of experimentally validated short peptides that undergo self-assembly to form ordered nanostructures manually curated and compiled from 301 research articles published in the recent past with the rise of interest in nanobiotechnology (Figure 3C). Of these 1,049 entries, 701 entries are of dipeptides, 328 entries belong to tripeptides, and 20 entries belong to single amino acid with their conjugate partners. 48 of the peptides were cyclic, while the rest were linear. These model short peptides form 13 major types of nanostructures, as shown in Figure 3A. Hydrogel (297 entries) is the most common type of structure followed by nanofibers (127 entries), nanotubes (67 entries), nanospheres (50 entries), nanoparticles (47), and hundreds of entries of mix nanostructures. Different techniques have been described in the literature to study the phenomenon of self-assembly of peptides. The most common techniques were scanning electron microscopy, transmission electron microscopy, atomic force microscopy, Fourier transform infrared spectroscopy (Figure 3D).

**Figure 3:**
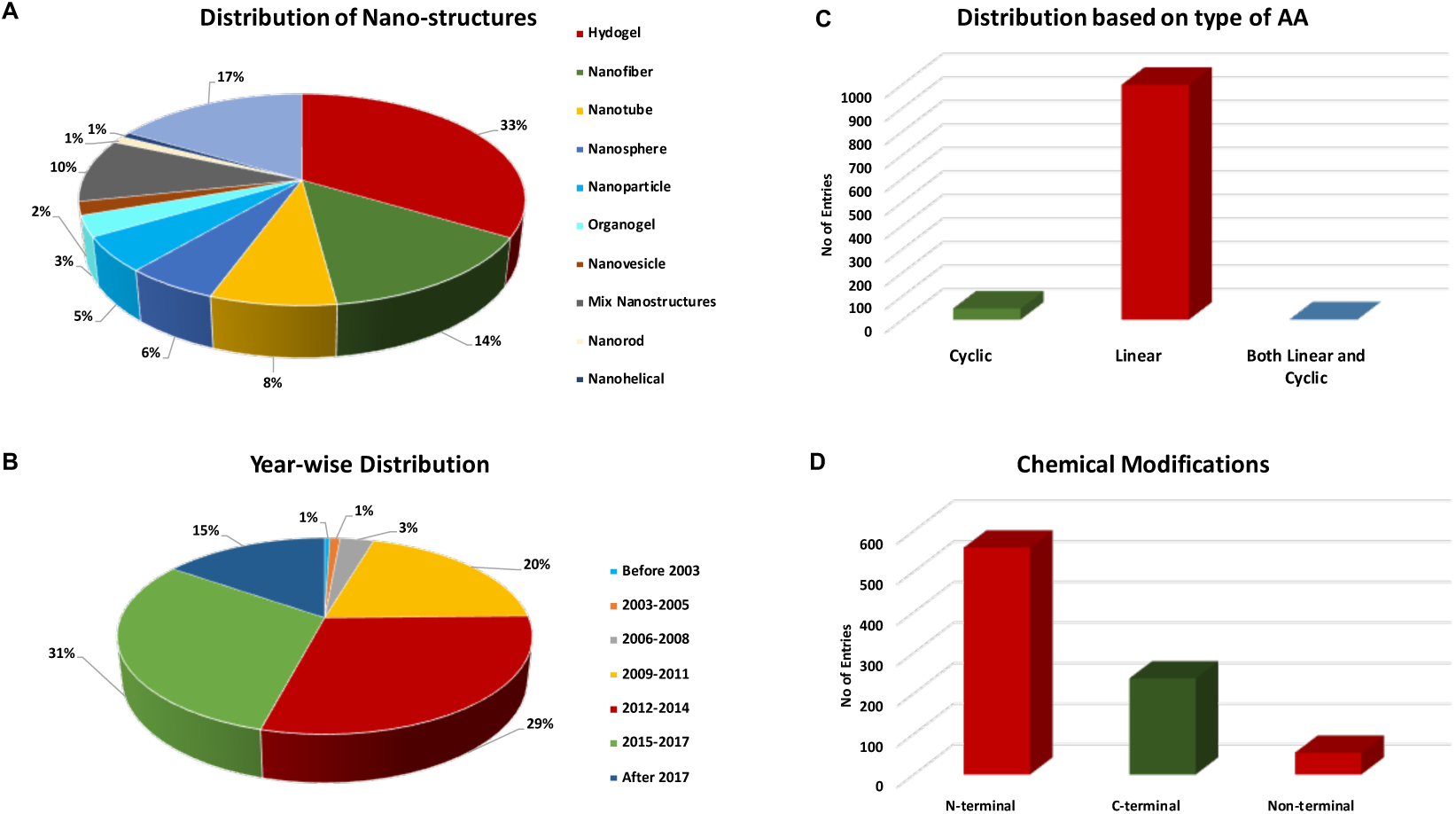
Statistics of distribution of entries in the SAPdb: (A) Types of nanostructures, (B) Year-wise distribution, (C) Based on Peptide, and (D) terminal modification.

To give different functionality, stability, and shape to nanostructure, some peptides have been cyclized. In contrast, various terminal modifications and non-natural amino acids have been incorporated in various other peptide sequences (Figure 3B). Thereis a total of 382 entries in SAPdb encompass modifications at the N-terminal. Most commonly reported N-terminal modifications include Fmoc (fluorenylmethoxycarbonyl; 160 entries), Boc (N-t-Butoxycarbonyl; 113 entries) and acetylation (46 entries); while, methoxylation (73), amidation (49 entries), benzylation (33 entries), were reported as primary modifications at C-terminal. Besides this, 54 reported entries had non-natural amino acids like dehydrophenylalanine (ΔPhe) (30 entries), beta-alanine (β-Ala) (16 entries), amino benzoic acid (ABA) (9 entries), amino isobutyric acid (Aib) (9 entries), *etc*. in their sequences.

### Implementation of Web Tools

Various modules, i.e., search tools, browse tools, and analysis tools, were integrated into the database for users to explore SAPdb.

### Search Tools

Two distinctive modules are implemented in SAPdb, i.e., Simple search, Advance search under search tool to assist the user for amiable data retrieval.

### Simple search

This module offers the fundamental facility for data retrieval from the database. Here one can perform a keyword query by selecting any required field of SAPdb such as the type of self-assembly, technique, peptide sequence, etc. Besides, this module also permits the users to select anticipated fields to be displayed in the result.

### Advance search

To retrieve relevant information from SAPdb, the advance search module presents the facility for the user to implement multiple query system. By default, it executes four queries simultaneously, but the user can select the desired keyword search from any selected field. Moreover, this module allows the implementation of standard logical operators (=, >, <, and LIKE). Besides this, advance search permits the user to integrate the output of different queries by employing operators like ‘AND and OR’. Further, the user can also add or remove the queries to be executed.

### Browsing Tools

The browsing facility has been integrated to facilitate the user for data mining in systematic mode from SAPdb. In this module, the user can fetch information on peptides by browsing six different classes (i) Chemical modification, (ii) Type of Nanostructure, (iii) Size of Nanostructure, (iv) Technique, (v) Publication year, (vi) Dipeptides and (vii) Tripeptides.

The chemical modification field assists the user in retrieving data on peptides that have different chemical modifications at the N or C termini. For instance, Fmoc (fluorenylmethoxycarbonyl; 160 entries), Boc (N-t-Butoxycarbonyl; 113 entries), and acetylation (47 entries) are major modification at N-terminal; while methoxy (73), amidation (49 entries), benzylation (33 entries), were reported as primary modifications at C-terminal. Type of nanostructure category offers the user to fetch detailed information on peptides that form a particular nanostructure such as nanotube, nanosphere, nanofibers, hydrogel, etc. The size of the nanostructure category allows users to browse entries based on the size of the nanostructure reported. Further technique category permits the user to extract information regarding peptides whose nanostructure was studied using a particular technique such like Transmission Electron Microscopy (TEM), Scanning Electron Microscopy (SEM), and Atomic Force Microscopy (AFM), and Fourier Transform Infrared (FTIR), etc. Additionally, users can also retrieve information of the peptides by the year of publication. Different entries of dipeptide and tripeptide sequences can be explored under the dipeptide and tripeptide modules of the browse facility.

### Analysis Tools

This module enables the users to execute several examinations, such as sequence similarity, peptide mapping, and physicochemical properties of peptides.

### GGSearch

This tool allows the user to perform an efficient similarity-based search against the short peptides in SAPdb. It is based on Needleman-Wunsch alignment [66].

### Smith–Waterman Algorithm

This algorithm [66] permits the user to search peptides in the SAPdb database similar to their query peptides. Under this option, the user can submit concurrently multiple peptide sequences in FASTA format.

### Peptide Mapping

This tool offers the facility to map SAPdb peptides over the query protein sequence. It will be useful for identifying motifs within proteins that tend to undergo self-assembly.

### Physicochemical Properties

This module facilitates users to examine properties like charge, polarity, volume, hydrophobicity, etc. of the desired peptide sequences.

### Effect of Experimental Conditions

Previous studies have shown that the variation in experimental conditions like the type of solvent, change in concentration of peptide, temperature, incubation time, pH, etc. can lead to the formation of different nanostructures by the same peptide [67]. Further, our analysis shows that the concentration of peptide is also an important factor for controlling the shape and size of the nanostructure. For example, KFG tripeptide forms vesicles at a low concentration of 0.5mg/ml while at higher concentration of 5mg/ml forms nanotubes [67]. Furthermore, the role of solvent in directing the type of nanostructure formation is also revealed through this analysis. Most of the nanostructures are formed using water as a solvent, followed by phosphate buffer and organic solvents like methanol, ethanol, hexafluoropropanol, chloroform, and acetone. FF is reported to form nanovesicles in acetone, while in water, it forms nanotubes [68]. This analysis also shows some specific experimental conditions favor the formation of nanostructure as evidenced by the observation that the majority of the nanostructures are reported to be formed by self-assembly under room temperature or 25 °C (499 entries) and in the range of neutral pH of 7 to physiological pH of 7.4 (178 entries) followed by acidic pH of less than 6 (118). Besides, we have observed that there 258 entries (see Table 1), where, peptides form different nanostructures without any chemical modifications or conjugate partners. Out of 258 entries, 180 (70%) entries contain any of aromatics amino acids. This indicates presence of aromatics amino acids favored the self-assembly formation.

### Case Study

Our analysis also shown that the different experimental condition changes the nature of self-assemblies. In order to understand the effect of conditions, we have considered an example of diphenylalanine (FF). FF forms hydrogels at different experimental condition such as temperature = 55°C, pH= ∼3-4, 7.2 and 8, incubation time >24 hours etc. On the other side, it forms nanostructure like nanosphere, nanotubes, and nanofibers at various conditions like pH, N-terminal modification, temperature, and incubation time, as represented in **Figure 4**.

**Figure 4:**
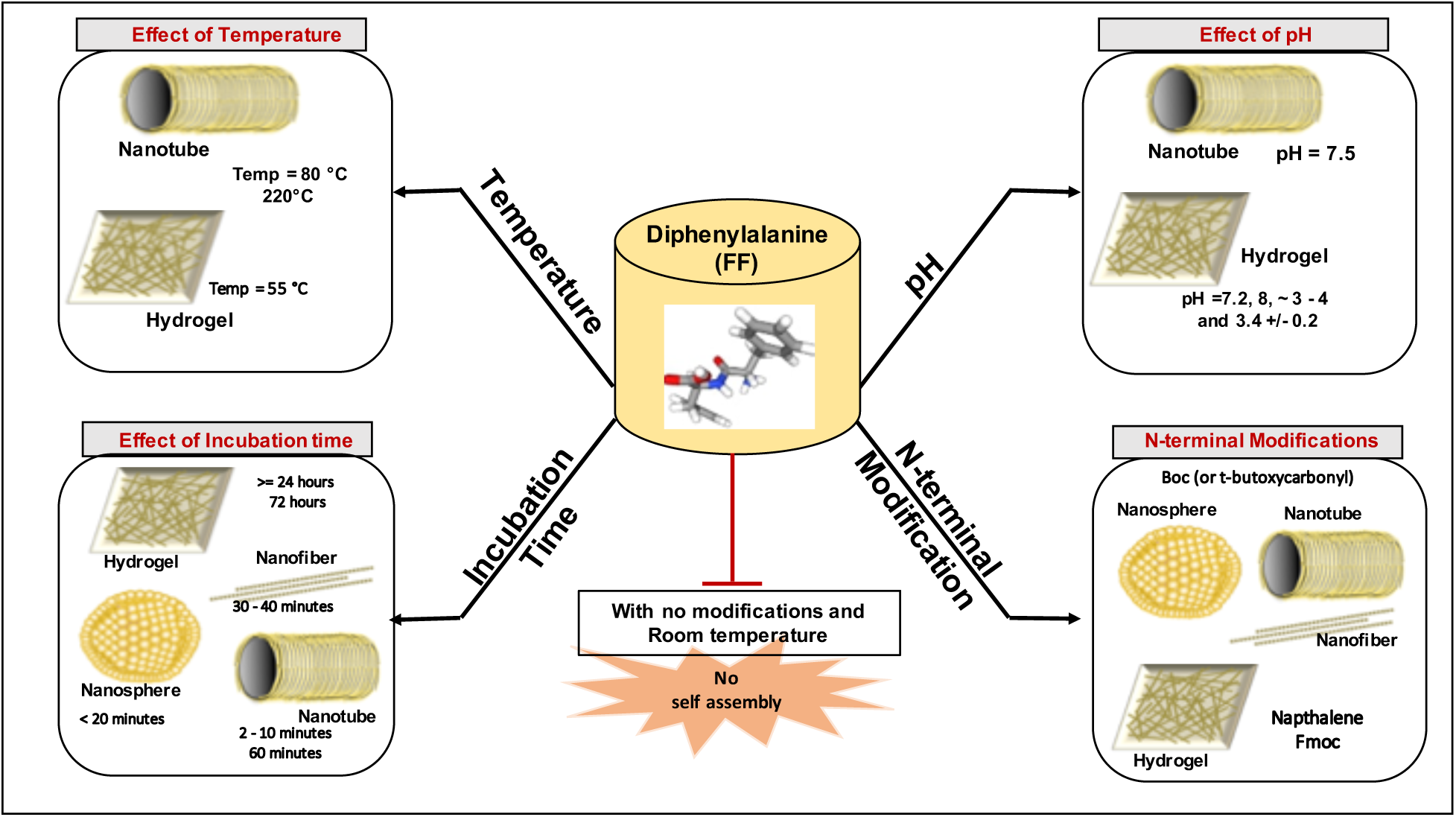
The demonstration of the effect of experimental conditions (pH, temperature, incubation time, and N-terminal modification) in the formation of different self-assembled structures like nanosphere, hydrogel, nanofiber, and nanotube.

In the absence of any user-friendly web-platform, the user needs to go through widely spread literature in the form of the bulky text publications. SAPdb allows easy access to such information in tabulated form, which leads to a better understanding of the 3D shape of nanostructures formed by altering experimental conditions.

### Working of SAPdb database

To understand how a user can be queried and retrieve data from the SAPdb database, we have demonstrated (Figure 5) with an example of a dipeptide, i.e., dipleneyalanine (FF). Figure 5 representing the step by step information on how users can query/search SAPdb using a simple search module by a keyword like PMID, Peptide sequence, Peptide name, year of publication, etc. For instance, here, we queried the search module with peptide sequence, i.e., FF in the given search space (Figure 5A) of the database. User can further select various display fields that are given on a simple search page. On submitting the desired query (e.g., FF) using the “Submit” button, the next page will display all the entries related to the query; for instance, here we retrieved 362 entries of FF, as shown in Figure 5B. Each entry has a unique SAPdb ID, and the user can get the detailed information regarding peptide by further clicking on the SAPdb ID. The next page provides complete information about the selected SAPdb ID, as represented in Figure 5C. It gives the peptide’s primary information (SAPdb ID, PMID, Year, Name, Sequence, N-terminal modification, C-terminal modification, non-terminal modification, peptide/conjugate/Mixture, conjugate partner, Technique, solvent, Method, concentration, pH, Temperature, etc. The secondary information provides Physiochemical properties, Structure, and SMILS of the given peptide. Users can further explore these properties, as shown in Figure 5D.

**Figure 5:**
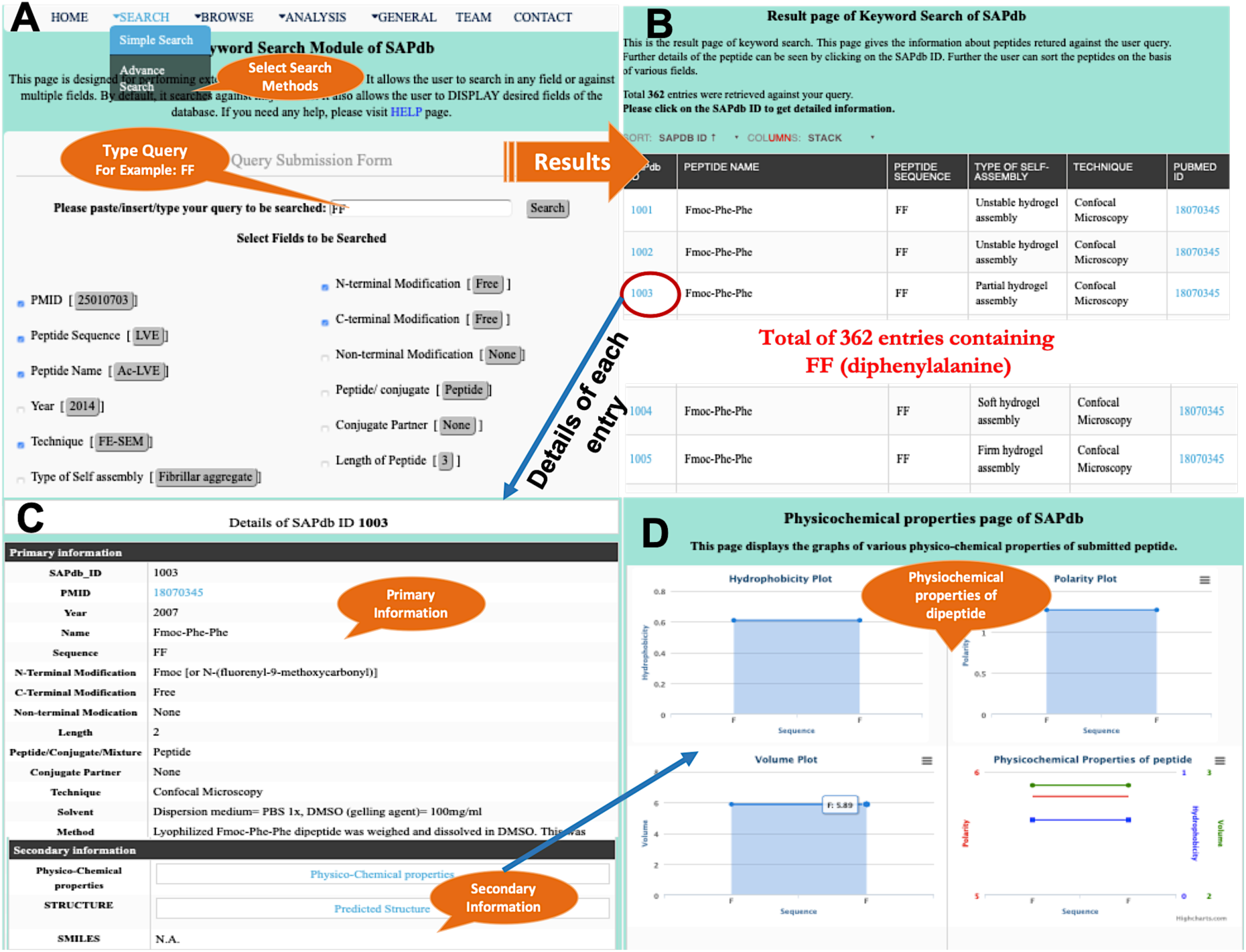
Demonstration of querying and retrieval of information from SAPdb using the Simple Search module.

## Discussion

With the rising interest in nanobiotechnology, it is important to understand the properties governing the self-assembly of peptides into nanostructures. Previously databases like AmyLoad [69], CPAD [70] and AMYPdb [71] have curated protein and peptide aggregation. Since there was no repository of experimentally validated and well-ordered self-assembling peptide nanostructures, we have developed SAPdb, a platform with comprehensive information about nanostructure formed by short peptides. Dipeptides and tripeptides are the shortest peptides that can assemble into higher-order structures. They are the model candidates to understand the process of self-assembly. SAPdb holds information of 167 unique dipeptides and 96 unique tripeptides that have been experimentally validated for undergoing ordered nanostructure formation by self-assembly.

Comprehensive analysis of SAPdb data have shown that different environmental conditions like temperature, pH, solvent, concentration, etc. direct the shape or type of nanostructures formation. Further, the specific conditions, i.e., room temperature, acidic pH, favors the formation of nanostructure of self-assembling peptides. Besides, amino acid composition analysis of peptides of SAPdb revealed that self-assembling peptides are rich in valine, tryptophan, leucine, and alanine. At the same time, they are depleted in isoleucine, aspartic acid, and histidine compared to non-self-assembling peptides. Further, aromatic residues like phenylalanine and tyrosine are favored at the second and third positions in tripeptides undergoing self-assembly. Besides, negatively charged residues are preferred at the C-terminal and while positively charged residue are at the N-terminal.

### Utilization of SAPdb

SAPdb maintains nanostructure-forming small peptides, which have numerous applications in diverse areas as illustrated by the following examples:

### The drug, gene, and other biomolecules delivery vehicles

One of the major applications of SAPs lies in the delivery of drugs and biomolecules. This is attributed to their biocompatible nature, the small size of self-assembling dipeptides and tripeptides [72–76]. For instance, EΔF dipeptide based nanovesicles employed to deliver vitamin B-12 in the HeLa cells [77], while MΔF, IΔF, and LΔF were used for the invitro delivery of curcumin in L-929 cells and in vivo delivery in mouse [78].

### Cell culture scaffold biomaterial for tissue engineering

Another major application of the peptide based self-assembled hydrogels is their efficiency for 3D-cell culture (Jayawarna). It has been observed that dipeptide FΔF based hydrogel sustained 3D cell growth of HeLa cells and L929 (mouse fibroblast) cells for more than 14 days, with substantial multiplication rate and cell viability [79].

### Models for nanofabrication and biomineralization

In the past, self-assembling peptides were explored as templates for nanofabrication like nanowires, biomineralization, nanocircuits owing to their potential of self-assembly [80, 81]. For example, Phe–Phe tagged with ferrocence forms uniform nanowires having a diameter of 100 nm with a length 5–10 mm. After adsorbing gold nanoparticles and antibody molecules onto these Fc-tagged nanowires surface, these nanowires were employed as a detection probe for sensitive electrochemical immunosensing [82].

We have incorporated peptides with such far-reaching applications in SAPdb so that the users can easily retrieve the SAPs of their interest. Using different modules, one can examine how the substitution of D-amino acids or non-natural amino acids in the peptide sequence and the experimental conditions like a solvent, concentration, pH, etc. influence the self-assembling property of the peptides. Moreover, to facilitate the users, we have integrated a chemical modification browsing tool to highlight the impact of different modifications on the type of nanostructure formed by the same peptide sequence. The users can easily analyze sequence similarity of their query peptide with different SAPs at a single platform, i.e., SAPdb, which are otherwise scattered in the literature and difficult to access.

### Future Developments

As the scientific community is actively publishing articles pertaining to this emerging field, in the future, our first concern will be to update data available about short model peptides as well as to expand the database further to include longer peptides which form ordered nanostructures on self-assembly.

## Acknowledgments

The authors are thankful to the Department of Science Technology, India (JC Bose National Fellowship) for financial support. D.M. and H.K. are grateful to the Council of Scientific and Industrial Research (CSIR), India, and A.D. and N.S. are thankful to the Department of Science Technology (DST), India, for providing fellowships.

## Author contributions

D.M., H.K., A.D., and N.S., collected and compiled the data. D.M., H.K., developed the website and penned the manuscript. G.P.S.R. conceived the idea and coordinated the project.

